# Parallel multiOMIC analysis reveals glutamine deprivation enhances directed differentiation of renal organoids

**DOI:** 10.1101/2025.02.27.640060

**Authors:** Iman Sarami, Katherine E. Hekman, Ashwani Kumar Gupta, Justin M. Snider, David Ivancic, Manja Zec, Manoj Kandpal, Issam Ben-Sahra, Rajasree Menon, Edgar A. Otto, Floyd H. Chilton, Jason A. Wertheim

## Abstract

Metabolic pathways play a critical role in driving differentiation but remain poorly understood in the development of kidney organoids. In this study, parallel metabolite and transcriptome profiling of differentiating human pluripotent stem cells (hPSCs) to multicellular renal organoids revealed key metabolic drivers of the differentiation process. In the early stage, transitioning from hPSCs to nephron progenitor cells (NPCs), both the glutamine and the alanine-aspartate-glutamate pathways changed significantly, as detected by enrichment and pathway impact analyses. Intriguingly, hPSCs maintained their ability to generate NPCs, even when deprived of both glutamine and glutamate. Surprisingly, single cell RNA-Seq analysis detected enhanced maturation and enrichment for podocytes under glutamine-deprived conditions. Together, these findings illustrate a novel role of glutamine metabolism in regulating podocyte development.

## Introduction

Renal cell types derived from human pluripotent stem cells (hPSCs) may one-day enable kidney disease modeling or cellular therapies that either restore or replace organ function. Self-organized, stem cell-derived renal organoid tissue has opened the door for development of patient-specific organ and tissue engineering ^1^. Renal organoids are also a unique tool to study the kidney micro-environment on both the molecular and cellular scale, to tease apart critical pathways in kidney development or to investigate causes of disease initiation in congenital abnormalities ^2–8^.

Several strategies have emerged to differentiate hPSCs toward renal organoids ^5,6,8–11^. These methods exhibit some variation, but all share common themes of differentiating hPSCs toward mesoderm and subsequently nephron progenitor cells (NPCs). After reaching the NPC stage, individual cell types self-organize into recognizable early renal structures within organoids. Despite significant advances, this process suffers from limited understanding of key genetic, epigenetic and metabolic pathways that control the differentiation process.

Metabolic reprogramming in stem cells, which are the metabolic changes during normal development or *in vitro* directed differentiation, provides important leads to better understand critical drivers of cellular differentiation^12^. In this study, we hypothesized that metabolic reprograming occurs during directed differentiation of hPSCs toward NPCs and subsequently toward renal organoids. Furthermore, these specific changes in metabolic pathways may be harnessed to influence the cellular phenotype within organoids and improve the differentiation process.

## Results

### Global metabolic profiling during differentiation of H9 to NPCs indicates significant changes in the alanine-aspartate-glutamate pathway

To optimize metabolic analyses, we generated renal organoids from the H9 human embryonic stem cell line (hESC, NIH Approval Number NIHhESC-10-0062) according to a modified version of the protocol by Morizane et al. (Figure S1A) ^10^. Differentiation followed the expected progression, with loss of *OCT4* expression and upregulation of NPC genes (*SIX2*, *PAX8*, *LHX1*, and *GATA3*) at D9 (Figure S1B, S1C). These NPCs self-organized into organoids bearing markers of podocytes (podocalyxin), proximal tubule cells (LTL) and distal tubular cells (E-cadherin) in both the 2D condition (data not shown) and 3D spheroid culture, by IHC (Figure S1D) and by TEM (Figure S1E). These findings validate our modified feeder– and serum-free approach generating renal organoids.

We then performed intracellular metabolomics analysis at days 0, 1, 3, 4, 6, 7, and 9, representing the beginning and the end of the major stages (Figure 1A) ^10^. The principal component analysis (PCA) revealed distinct metabolic profiles in pluripotency (D0), mesoderm (D3), intermediate mesoderm (D7) and the NPC stage (D9) (Figure1B). Individual metabolites were ordered according to hierarchical clustering and revealed three main patterns: An increasing trajectory, such as taurine and myo-inositol; a decreasing trajectory, such as glutamine and glutamate; and a mid-differentiation maximum (e.g., an inverse U-shape pattern), such as cytidine (Figure 1C). Enrichment analysis and pathway impact scores revealed significant changes in multiple pathways (Figure 1D, 1E). Among the decreasing metabolites, the alanine, aspartate and glutamate pathway changed significantly, by both enrichment and pathway impact scores. Individual metabolites within the alanine, aspartate and glutamate pathway demonstrated a decreasing pattern after D3 of differentiation, with only two exceptions, alanine and carbamoyl-phosphate (Figure 2A-B). The altered carbamoyl-phosphate indicated increased metabolism into the urea cycle, and while the urea cycle metabolites ornithine, citrulline and urea demonstrated increases during differentiation, the ratio of intracellular/extracellular urea remained relatively stable (18%), indicating the increased urea was cleared from the cell, potentially avoiding toxic levels of ammonia (Figure 2C). Intracellular lactate increased early during differentiation and demonstrated a negative correlation with multiple lipid oxidation metabolites, suggesting an increase in cytosolic glycolysis, until D6, and subsequently decreased toward the NPC stage (Figures 2D), which further supports a change in energy utilization and oxidative phosphorylation, i.e., metabolic reprogramming ^13–16^. These findings illustrate major changes in the metabolic profile during differentiation to NPCs, notably in the alanine, aspartate and glutamate pathway.

**Figure 1:**
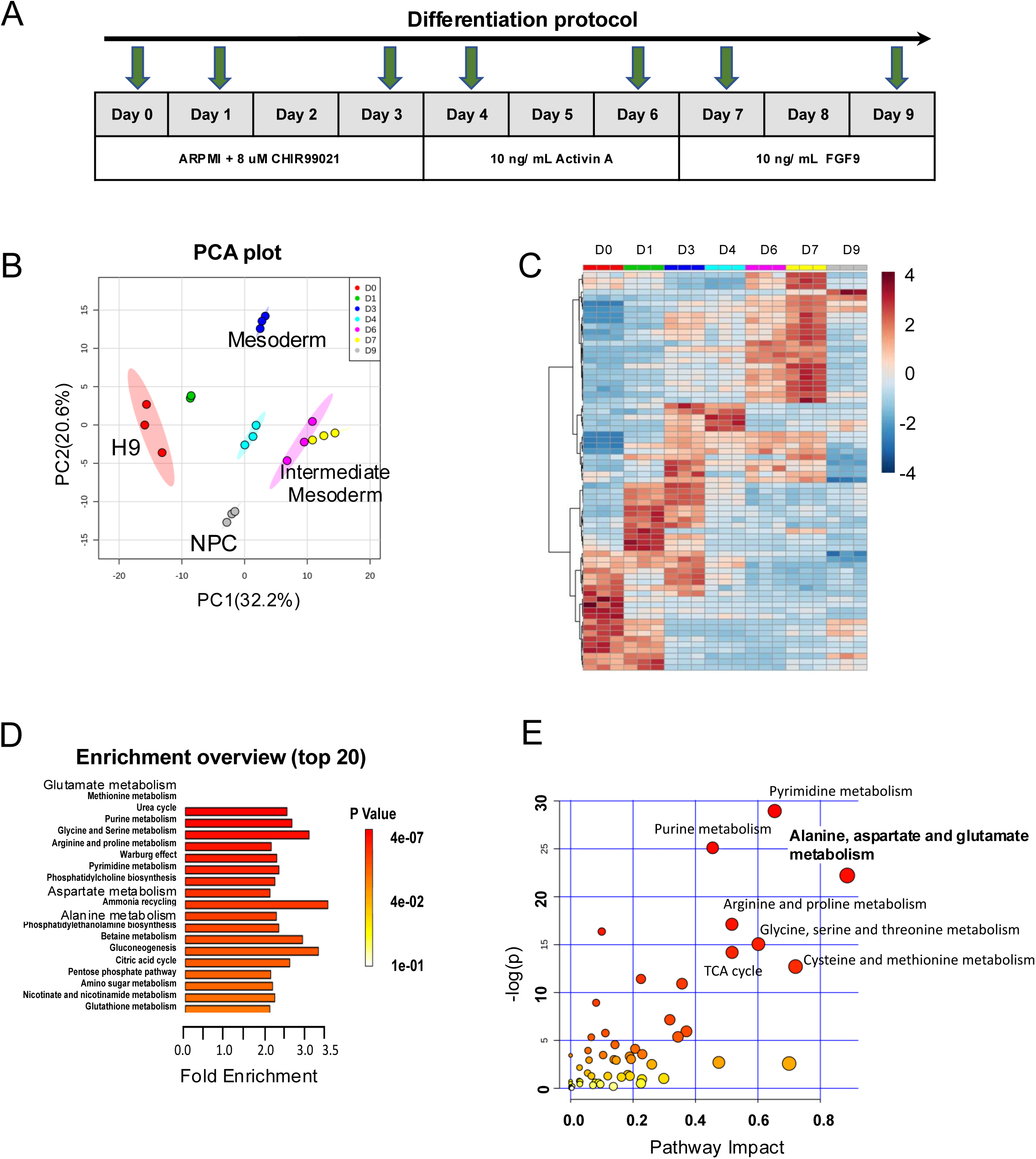
*Intracellular metabolomics analysis of H9 during differentiation into NPCs.* A) Differentiation protocol from pluripotency (Day 0) to NPCs (Day 9). Green arrows indicate time-points for sample collection. B) PCA analysis reveals distinct metabolic profiles from pluripotency (D0) to NPCs (D9). Individual dots represent biological replicates, N=3. C) Heatmap showing metabolite concentrations (red indicates high, blue indicates low) following three main patterns: Increasing, decreasing and a mid-term maximum at the middle of the differentiation process. D) Pathway enrichment analysis of identified metabolites that significantly change over the early differentiation period to NPCs. E) The pathway impact analysis of metabolites using MetaboAnalyst 4.0 identifies the alanine, aspartate and glutamate pathway as having the highest impact.

**Figure 2:**
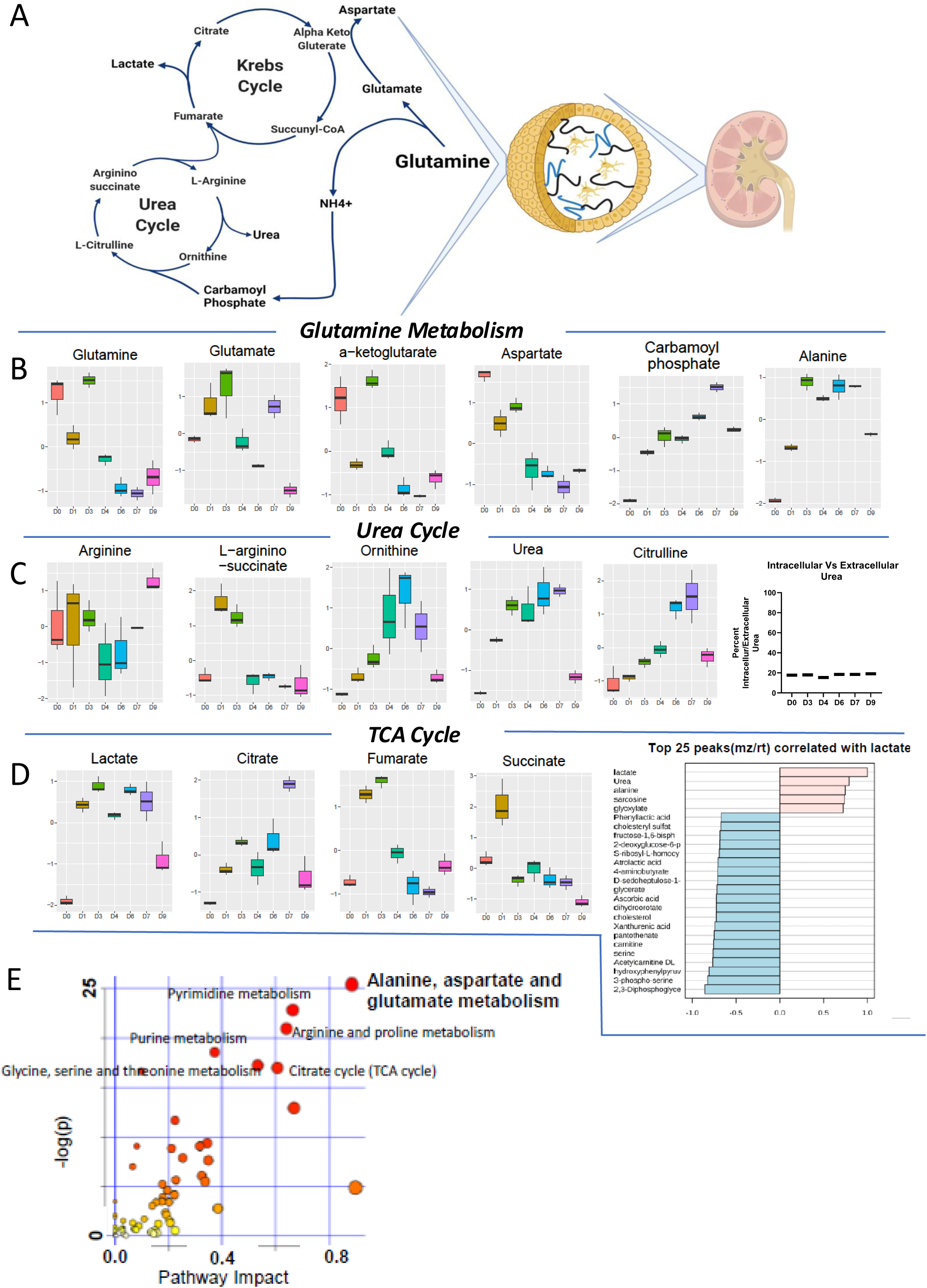
*Intra– and extracellular metabolomics correlate during H9 differentiation to NPCs.* A) Overview of glutamine metabolism. B) Changes in intracellular concentration of alanine, aspartate, and glutamate pathway metabolites from D0 to D9. C) Changes in urea cycle metabolites from D0 to D9. D) Changes in lactate metabolism (left) and top 25 metabolites that correlate with intracellular lactate concentration (right). E) Pathway impact analysis of metabolites using MetaboAnalyst 4.0 identifies the alanine, aspartate and glutamate as having the highest pathway impact by extracellular metabolomic analysis.

We next performed extracellular metabolomics analysis during differentiation (D0 – collected 1 hour after commencing differentiation, D2, D3, D4, D5, D6, D7, D8, D9) to better understand environmental effectors and intercellular signaling during maturation. Similar to intracellular metabolomics, distinct metabolic profiles were found at the pluripotent, mesoderm and NPC stages by PCA (Figure S2A). The extracellular heatmap data likewise revealed the same three patterns as the intracellular metabolomics (Figure S2B). The decreasing pattern was comprised primarily of media components taken up by cells during differentiation, such as cystine and serine (Figure S2C). The levels of extracellular metabolites generally correlated inversely with the corresponding intracellular pattern. These data indicate that the increased energy burden of differentiating cells was met by increased uptake of metabolites from the media.

Enrichment and pathway analyses of extracellular metabolite levels revealed the pathways most significantly altered during differentiation which also included the alanine, aspartate and glutamate pathway (Figure S2D, 2E), further supporting the importance of this pathway in the differentiation to NPCs.

### Human induced pluripotent stem cells (hiPSCs) undergo similar metabolic shifts during differentiation to NPCs and renal organoids

While both hESCs and human hiPSCs share self-renewal and pluripotency properties, they each may be influenced by different epigenetic drivers ^17^. To evaluate metabolic trends of differentiating hiPSCs to renal organoids, we performed intracellular metabolomics analysis at each stage of differentiation, D0, D1, D3 (mesoderm), D6 (intermediate mesoderm), D9 (NPC), D17 (pre-organoid) and D23 (organoid) (Figure 3A). As noted with differentiation of H9 cells, hierarchical clustering indicated large changes in metabolic profiles early in differentiation to the NPC stage (Figure 3B). PCA confirmed distinct overall metabolic profiles between stages (Figure 3C). Furthermore, the hiPSC (D0) stage demonstrated a metabolic profile distinct from NPCs (D9), pre-organoid (D17) and organoid (D23) stages, but these later stages shared similar metabolic patterns (Figure 3C). While the NPC (D9) and organoid (D23) stages were largely similar, the volcano plot did show a statistically significant decrease in glutathione and D-gluconate from NPC to organoid stage (Figure 3D). A direct pairwise comparison was performed on metabolomic data between D0 (hESC state) and D9 (NPC state), confirming distinct metabolic profiles by PCA plot (Figure 3E). However, continuity of metabolic programming of hPSCs toward organoids is depicted in a PCA plot (Figure 3F). This chart demonstrates metabolic profiles during days 1-9 for both hPSC cell types, which cluster together at each day, regardless of the cell type of origin. These data demonstrate the patterns of metabolic reprogramming during formation of organoids supersedes cell type.

**Figure 3:**
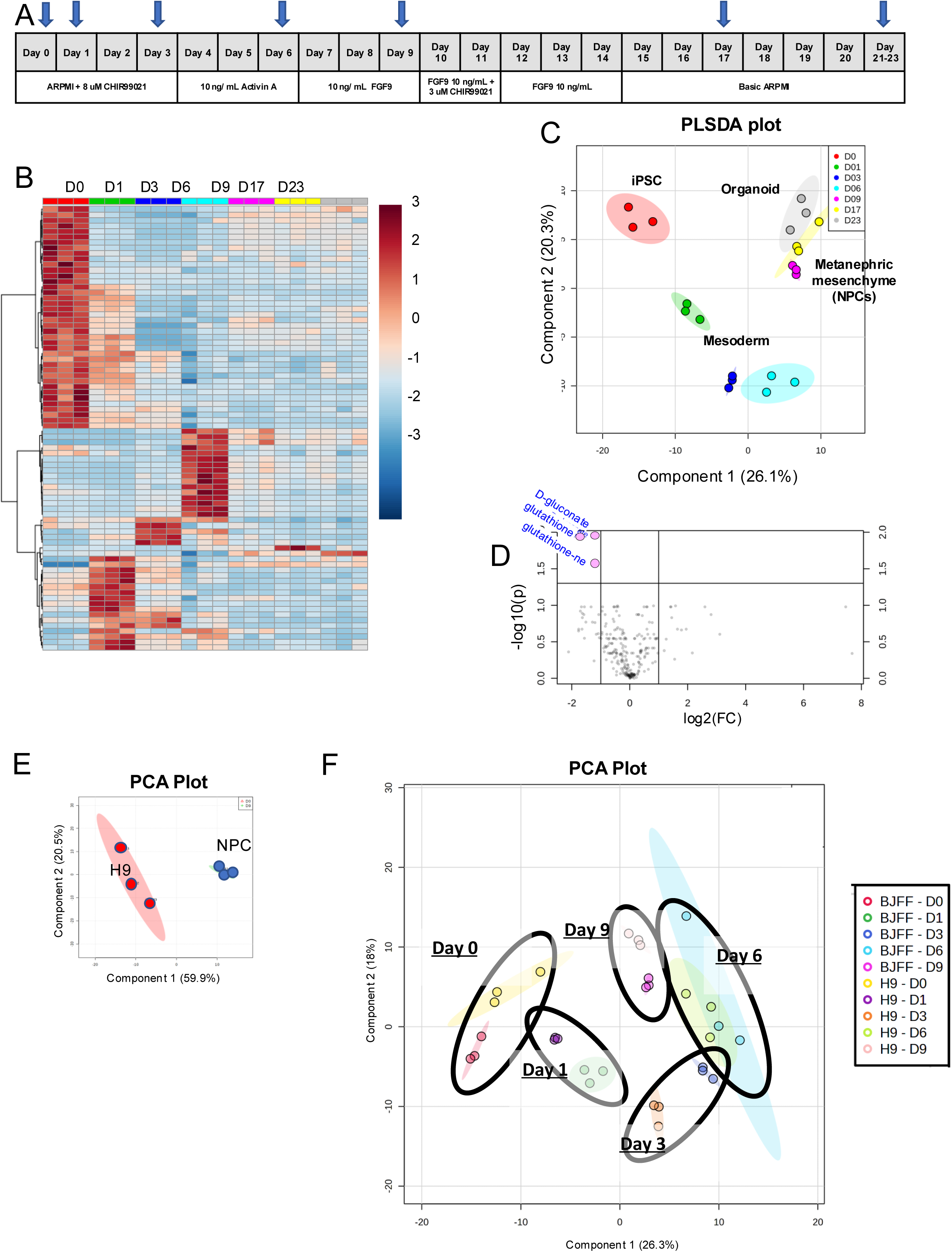
*Intracellular metabolomics analysis of hiPSCs during differentiation to NPCs and renal organoids and joint transcriptome-metabolome analysis of H9 differentiation to NPCs and renal organoids.* A) Differentiation protocol of hiPSCs to NPCs and renal organoids, blue arrows depict timepoints for sample collection. B) Heatmap represents metabolite concentrations following three main patterns (red indicates high, blue indicates low): increasing, decreasing and a mid-term maximum in the middle of the differentiation process. N=3. C) PCA analysis reveals distinct metabolic profiles between hiPSCs (D0) and NPCs (D9); however, NPC (D9, purple) and organoid stages show related profiles. Individual dots represent biological replicates. D) Volcano plot showing that only glutathione and gluconate undergo significant expression changes between NPC and organoid stage. E) PCA analysis reveals distinct metabolic profiles between D0 (pluripotency stage) and D9 (NPC stage). Individual dots represent biological replicates. F) PCA analysis of metabolic profiles for hiPSCs and hESCs on day 1-9, identical timepoints are circled to demonstrate clusters. Individual dots represent biological replicates.

### Integrated transcriptome and metabolome profiles during differentiation to NPCs demonstrate significant alterations in the alanine-aspartate-glutamate pathway

To further evaluate these changes in metabolic profiles, three replicates of whole-transcriptome sequencing (RNA-Seq) were carried out during differentiation of H9 cells to NPCs at the same seven timepoints used for the metabolomic analysis. Pathway and enrichment analyses showed that arginine and proline metabolism, pyrimidine metabolism and alanine, aspartate and glutamate metabolism were the most significantly altered pathways between these two timepoints (Figure 4A-B).

**Figure 4:**
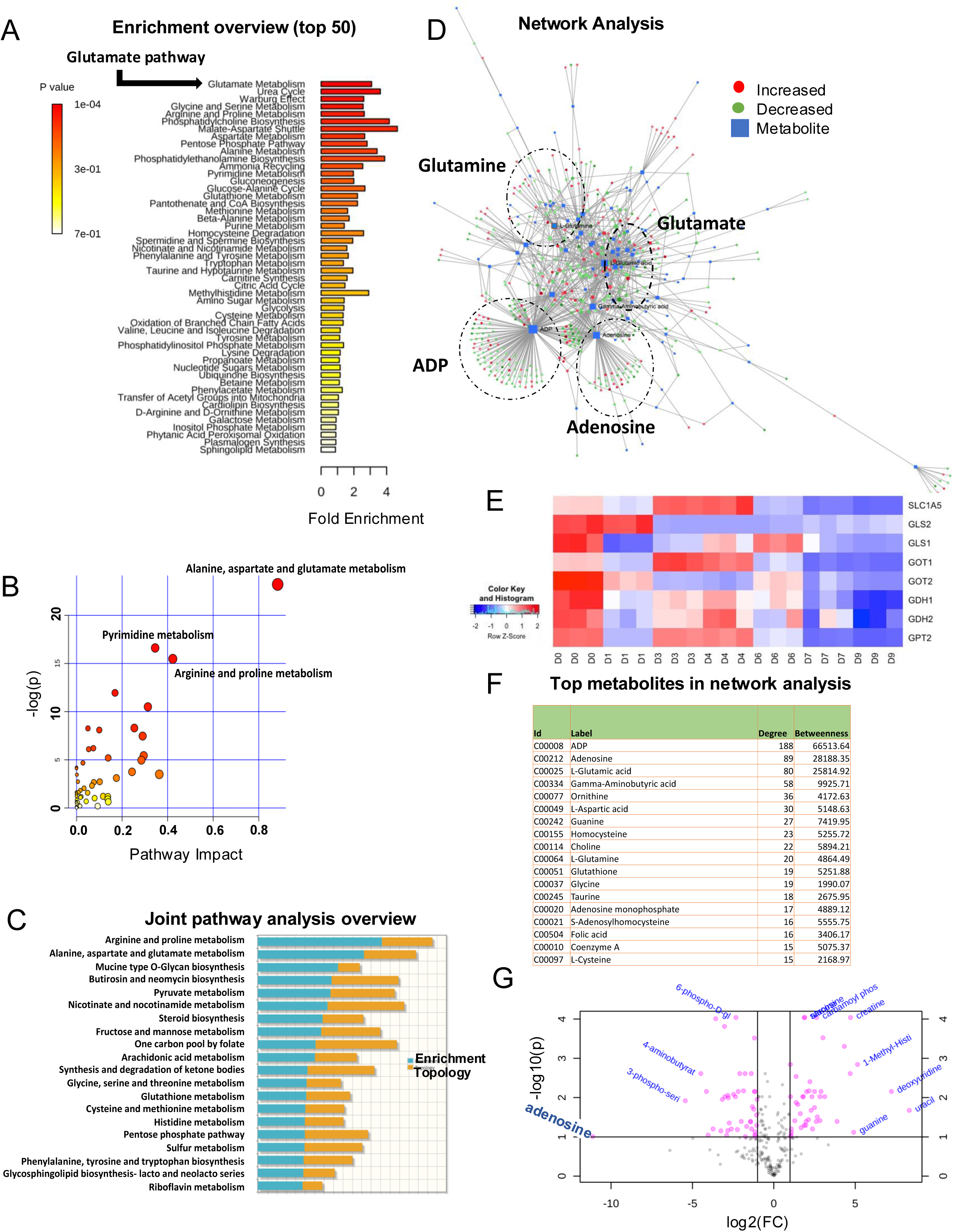
*Joint RNA-Seq and metabolomic analysis between H9 cells and NPCs.* A) Enrichment analysis reveals glutamate as a top altered pathway. B) Pathway impact scores of hESCs versus NPCs shows the alanine, aspartate and glutamate pathway with the highest impact. C) Joint pathway analysis between intracellular metabolomics and RNA-Seq between hESC and NPC stages (D0 vs D9). D) Network analysis showing the relationship between top metabolites and gene expression (D0 vs D9). E) Glutamine-related gene expression decreases over the course of NPC differentiation. F) Top metabolites found in network analysis. G) Volcano plot demonstrates a 100-fold decrease in adenosine between D9 and D0.

A joint transcriptome-metabolome pathway analysis was performed between significantly altered metabolites (FC>1.5) and genes (FC>2), and glutamate, using MetaboAnalyst 4.0 software (web based, first accessed March 2018) ^18^. In the joint analysis, arginine and proline metabolism (p-value = 0.008) and alanine, aspartate and glutamate metabolism (p-value = 0.019) were the top-most significantly altered pathways, with topology scores 1.0323 and 1.0612, respectively (Figure 4C).

Metabolic and transcriptomic data were further analyzed using a network analysis of significantly altered genes (FC>2) and significantly altered metabolites to determine the alteration of gene expression of each metabolite, consistent with the joint pathway analysis (Figure 4D). Furthermore, expression of glutamine-related genes decreased over the course of differentiation (Figure 4E). Among the top altered metabolites (Figure 4F), the intracellular adenosine concentration demonstrated the most significant change by volcano plot between D9 and D0, decreasing more than 100-fold (Figure 4G). This further supports the change in energy utilization between pluripotent H9 cells and NPCs. Together, the joint pathway and network analyses further support the major changes in metabolic profile of hESCs differentiating to NPCs, and highlight the importance of the alanine, aspartate and glutamate pathway.

### Only certain amino acids are essential for differentiation toward NPCs

Metabolomics and sequencing analysis highlighted specific pathways that each play critical roles in differentiation. To test the role of each amino acid in differentiation, we systematically removed each amino acid component, one at a time, from the media formulation and performed directed differentiation of H9 cells to NPCs. Deprivation conditions for arginine, cystine, histidine, isoleucine, leucine, lysine, methionine, phenylalanine, serine, threonine, tryptophan, tyrosine or valine were incompatible with cell survival from pluripotency at day 0 to day 3 of the differentiation process (data not shown). The remaining conditions allowed for cell survival through D9. These latter conditions displayed expected morphologic development (Figure S3A) with expression of SIX2 (Figure S3B), as well as some PAX8 (Figure S3C), by immunofluorescence.

These results are largely consistent with the extracellular metabolomic analysis and amino acid uptake. Specifically, extracellular metabolomics revealed a high uptake of arginine, cystine, threonine, tryptophan, valine, methionine, leucine, isoleucine, histidine and phenylalanine in the first 48 hours. In addition, alanine, asparagine, proline and glutamate concentrations in the media increased during differentiation, suggesting excretion of these amino acids. However, despite high uptake of glutamine and aspartate, H9 cells survived deprivation, suggesting a role for these metabolites beyond cell survival.

### Total glutamine deprivation enhances PAX8 expression during NPC generation

We hypothesized that glutamine plays a crucial role in NPC maturation. To test this hypothesis, we performed directed differentiation of H9 cells to NPCs in the presence and absence of glutamine. Deprivation of glutamine during differentiation to NPCs led to an earlier appearance of NPC-like morphology with continued self-organization into renal organoids (Figure S4A); however, glutamine deprivation led to a lower number of total NPCs by ∼50%, in comparison to NPCs derived in the presence of glutamine (Figure S4B-D). This was not due to increased cell death (Figure S4E). Glutamine deprivation also led to differential gene expression at D9 when compared to the complete media control, with 676 out of 21,794 genes being significantly altered in their expression with a >2-fold change (FC), 337 up-regulated and 339 down-regulated (Figure S4F). The advanced morphology was further augmented by combined deprivation of glutamine and glutamate (Figure 5A). We evaluated for key markers of NPC differentiation, SIX2 and PAX8, by immunofluorescence in NPCs generated under normal or glutamate and glutamine deprivation conditions. While all conditions expressed SIX2, expression of PAX8 varied (Figure 6B). The percent of PAX8 positive cells by flow cytometry at the end of D9 was the highest either without glutamine and glutamate (54.4 ± 2.1%) or without glutamine (47.1 ± 4.1%), which were statistically significant compared to the control condition (19.1 ± 2.0%) or without glutamate (21.1± 1.9%, p<0.0001, 2-sided type 2 student’s t-test, n=6 for D0 and n=9 for each NPC condition) (Figure 6C; sample gating strategy in Figure S5). These findings support a role for glutamine in regulating NPC differentiation.

**Figure 5:**
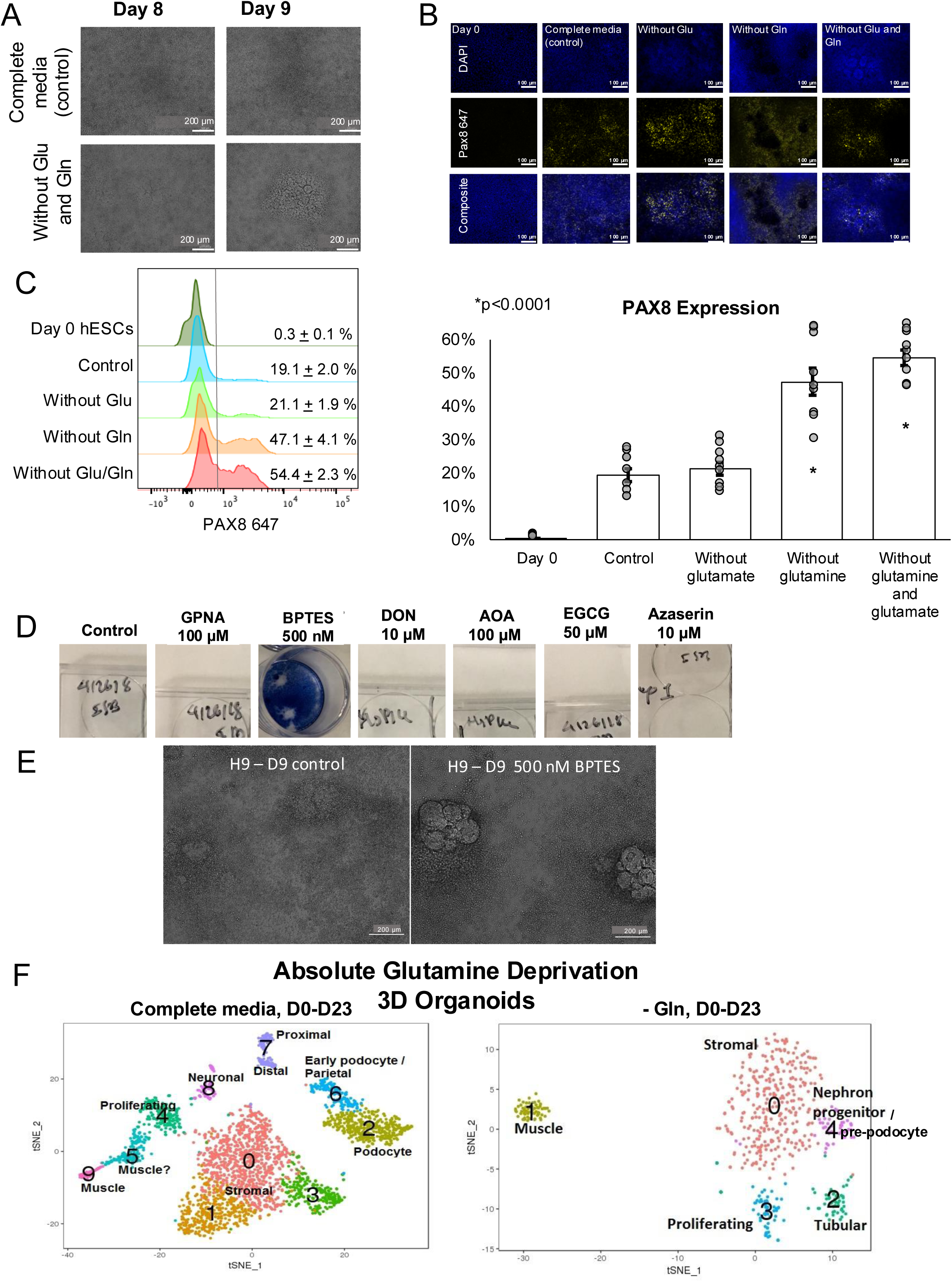
*Total glutamine deprivation accelerates PAX8 expression during differentiation to NPCs.* A) Deprivation of both glutamine and glutamate accelerated the acquisition of pre-tubular aggregate morphology by light microscopy. N=3. B) Immunofluorescence for PAX8 in the absence of glutamine, glutamate or both. Day 0, complete media, without Glu or Gln is reproduced from Figure S3C to illustrate a comparison to the absence of both Glu and Gln. Scale bar 100 µm. C) Glutamine deprivation significantly enhanced the number of PAX8 positive cells by flow cytometry at the end of D9. The bar graph illustrates the mean ± SEMs. All individual replicates are presented as gray dots. N=6 for control, N=9 for all others. D) The effect of glutamine metabolism inhibitors on the differentiation of H9 cells to NPCs. N=3. E) Representative image of nephron progenitor cells cultured in the presence of 500 nM BPTES demonstrate advanced morphology on day 9 comparable to being generated in the absence of glutamine. N=3 F) Droplet based scRNA-Seq of organoids generated in the presence (complete media) and total absence of glutamine in 3D culture.

### Entry of glutamine metabolites into the TCA cycle is essential for NPC generation

To further confirm the role of glutamine in the differentiation and maturation toward NPCs, we investigated the effect of inhibiting glutaminolysis on differentiation (Figure 5D). Upon reaching the NPC stage, Trypan blue was applied to the cells prior to desiccation to allow visualization of gross cell survival. Differentiating hESCs were intolerant to aminooxyacetate (AOA), a non-specific transaminase inhibitor that inhibits entry of glutamine metabolites into the TCA cycle ^19^. Inhibiting glutamine metabolizing enzymes with glutamine analogues azaserine and DON was also not tolerated, with no cells surviving to D9 ^20,21^. Inhibiting oxidative deamination of glutamate into α-ketoglutarate with epigallocatechin gallate (EGCG) during differentiation toward NPCs was also not tolerated. Conversely, inhibiting cellular uptake of L-glutamine with L-γ-Glutamyl-p-nitroanilide (GPNA) was partially tolerated ^22^. Bis-2-(5-phenylacetamido-1,3,4-thiadiazol-2-yl)ethyl sulfide (BPTES) is a selective inhibitor of kidney-type glutaminase isoform (GLS1), and prevents glutamine from entering the TCA cycle ^23^. BPTES was tolerated at 500 nM but not at higher concentrations and led to an NPC morphology mimicking glutamine deprivation (Figure 5E). Together, these data suggest that the glutamine deprivation phenotype can be reproduced pharmacologically, and that glutamine entering into the TCA cycle plays a key regulatory role in the differentiation of stem cells toward NPCs, while inhibition of glutamine metabolites in the cytoplasm or mitochondria is incompatible with cell survival to the NPC stage.

### Glutamine deprivation alters the course of differentiation in renal organoids shown by single cell RNA-Seq

To determine the cellular composition of 3D organoids developed under complete glutamine-deprivation and under standard conditions we performed single cell RNA-sequencing in both groups. This resulted in 543 total cell transcriptomes derived from the glutamine-deprived organoids and 2,799 transcriptomes which were generated from the organoids grown under the standard condition, after filtering out cells with less than 500 genes and more than 25% mitochondrial content. Unsupervised clustering was performed and a t-distributed stochastic neighbor embedding (t-SNE) plot for two-dimensional visualization was generated for each experimental group. Clusters were identified and labeled according to well-known differentially expressed marker genes. We identified ten major clusters of cells in organoids generated in the presence of glutamine including renal cells: cluster 2-podocytes, cluster 6-early/parietal podocytes, cluster 7-proximal and distal tubular cells; cluster 8-neuronal cells; clusters 5 and 9-two distinct population of muscle cells; clusters 0, 1 and 3-three distinct clusters of mesenchymal cells including renal mesenchymal cells; and cluster 4-proliferating cells (Figure 5F, left). Our findings are comparable with previous reports on single cell analysis of renal organoids derived from stem cells.^30^ In comparison, organoids derived from H9 cells in the absence of glutamine depict five distinct populations of cells including cluster 4-NPC/pre-podocyte and cluster 2-tubular cells; cluster 1-muscle cells; cluster 0-mesenchymal cells; and cluster 3-a population of proliferating cells (Figure 5F, right). In addition, we identified expression of *MEIS1* which is a marker of kidney stroma in one of the stromal clusters within organoids grown in the presence of glutamine. However, *MEIS1* was absent in stromal cells in glutamine-deprived organoids (Figure S6A). Overall, the data supports that glutamine deprivation alters the maturation and differentiation of H9 stem cells into renal organoids.

### Short-term glutamine deprivation during the NPC stage enhances podocyte maturation in renal organoids shown by single cell RNA-Seq

To distinguish whether the effect of glutamine deprivation is from selective survival in glutamine deprivation or a signaling effect of glutamine during differentiation, we deprived glutamine from differentiating cells for 24 hours at D8-9, a critical time point at which cells become nephron progenitor cells. We then interrogated gene expression with single cell RNA-Seq (DropSeq) to analyze all cell populations in both groups of resulting 2D organoids at D23. After filtering out cells with less than 500 genes and more than 25% mitochondrial content, single cell RNA-Seq resulted in 2,945 total cell transcriptomes derived from the glutamine-deprived organoids and 2,913 transcriptomes generated from the organoids grown under standard condition. Unsupervised clustering of all cell transcriptomes resulted in various clusters that were assigned to specific cell types/populations. Under standard conditions with glutamine (complete media control), 10 cell/transcriptome clusters were identified, similar to our previous results in the 3D organoids (heatmap shown in Figure S6B). In comparison, short term-glutamine deprivation yielded 11 clusters of cells (heatmap shown in Figure S6C). The datasets were combined via canonical correlation analysis (CCA) to permit direct comparison. Interestingly, two distinct podocyte populations were detected in the combined dataset (Figure 6A). Both podocyte populations contained cells from each condition (Figure 6B). The top 10 differentially expressed genes in each contained canonical podocyte markers, but each had unique features as well (Figure 6C). Interestingly, both podocyte clusters were predominantly populated by the glutamine deprivation condition (Figure 6D). Globally, podocyte markers, especially *NPHS2,* showed increased expression in the glutamine deprivation condition (Figure 6E, S6C). We validated our cluster annotation with the DevKidCC program, which confirmed our annotation of the two podocyte clusters, and both were predominantly populated by the glutamine deprivation population (Figures 6F-G). These data demonstrate that short-term glutamine deprivation at the NPC stage enhances the expression of mature podocyte markers in organoids.

**Figure 6:**
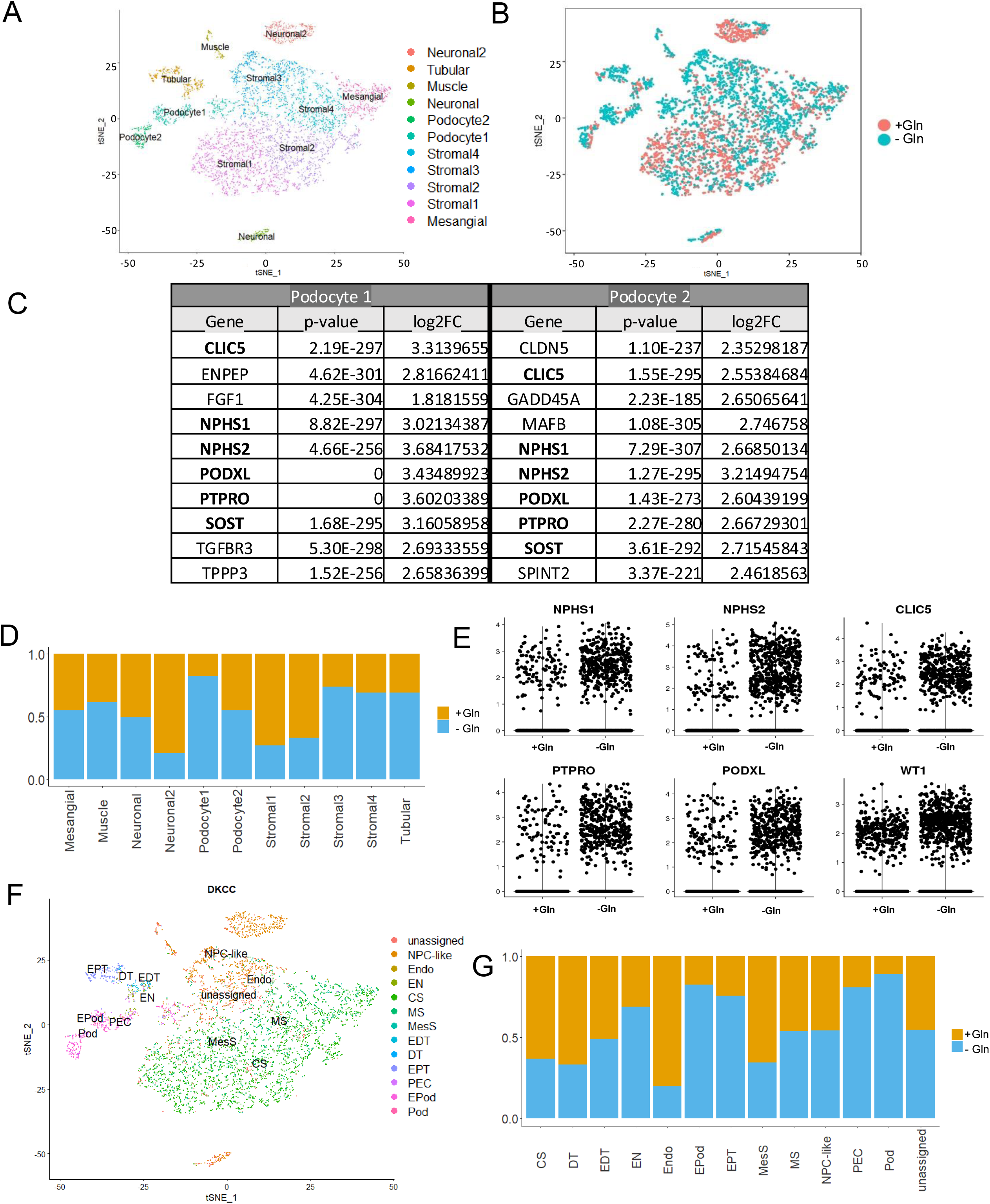
*Droplet based scRNA-Seq depicts the effect of short-term (24 hours) glutamine deprivation at the NPC stage on differentiation of hESCs to renal organoids in 2D culture.* A) Single cell RNAseq datasets of 2D organoids generated under control (complete medium; C1) and short-term glutamine deprivation (C2) were co-clustered via CCA for direct comparison. 11 clusters resulted, including 2 identified as podocytes by supervised cluster annotation. B) Red and blue highlighting of the C1 (+Gln) and C2 (-Gln) contributions, respectively, to each cluster. C) Top 10 differentially expressed genes in each of the two podocyte clusters. D) Relative contributions from C1 (+Gln) and C2 (-Gln) populations to each cluster, demonstrating the podocyte clusters are predominantly from C2 (short-term glutamine deprivation). E) Expression of podocyte markers in C1 (+Gln) and C2 (-Gln). F) DevKidCC cluster annotation confirms our supervised cluster annotation of podocyte clusters. G) Relative contributions from C1 (+Gln) and C2 (-Gln) to clusters as annotated by DevKidCC.

Next, we sought to characterize the impact of short-term glutamine deprivation on the development of podocytes. 2D and 3D organoids grown with or without short term glutamine deprivation were sectioned and stained positively for podocin, E-cadherin and LTL (Figure 7A). Podocin positive clusters in 3D organoids were counted in Fiji, revealing a significant increase in podocytes in the short-term glutamine deprivation condition. Together, these data suggest that glutamine-related signaling likely contributes to the underlying signaling mechanism triggering podocyte development.

**Figure 7:**
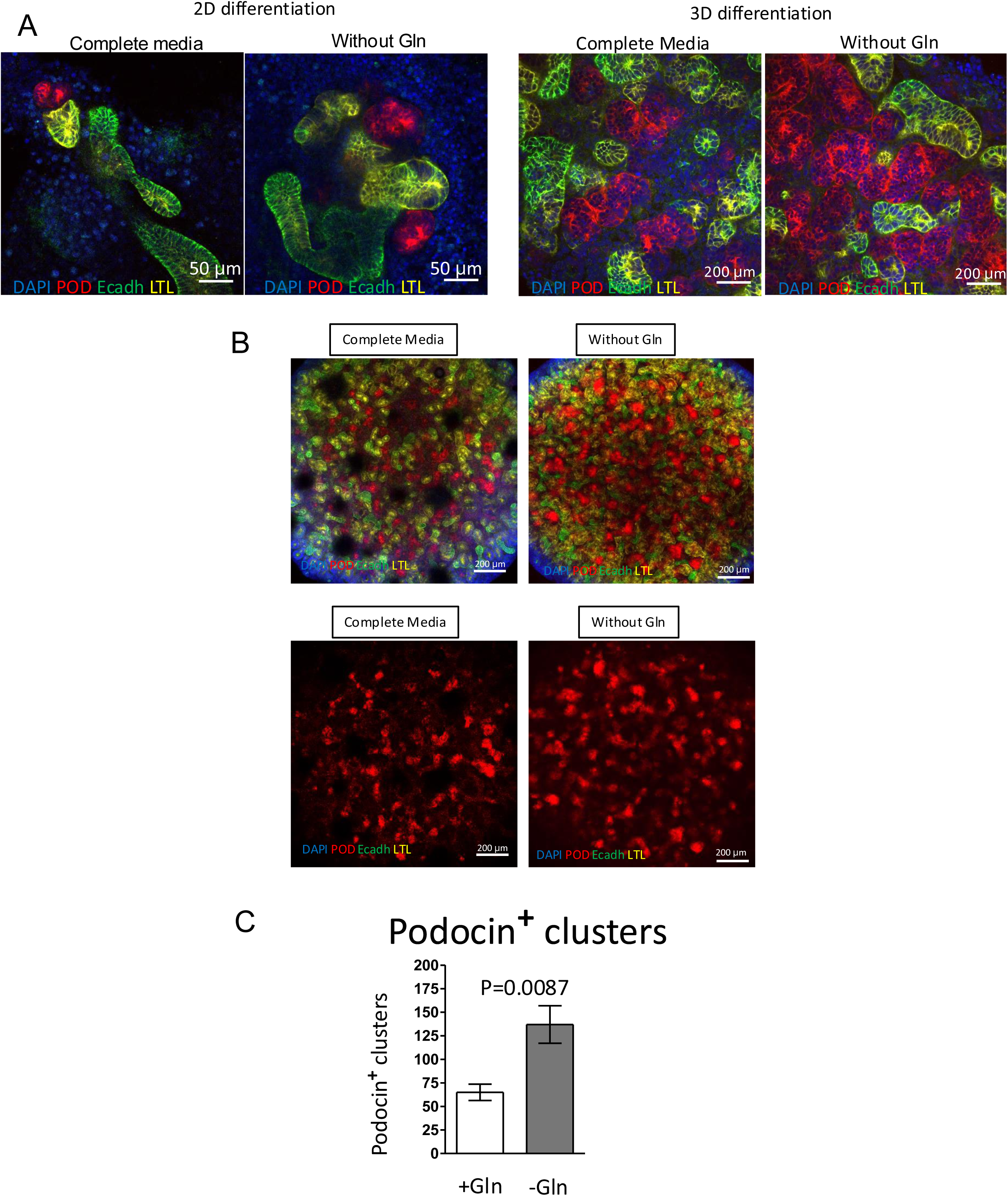
*Short-term glutamine deprivation enhances podocyte development.* A) Differentiation of H9 cells to 2D and 3D organoids with short-term glutamine deprivation yields staining for key kidney organoid components, with enhanced podocyte development. B) Staining of 3D organoids demonstrates increased podocin+ clusters in the short-term glutamine deprivation condition. C) Quantification of podocin+ clusters in Fiji. *p<0.05 by Student’s t-test. N=6

## Discussion

Our study is among the first to report parallel multiOMICS – a combination of single cell analysis, metabolomics and RNA sequencing – to investigate the underlying mechanisms of metabolic reprogramming in renal organoid development and stem cell differentiation. Our results demonstrate a critical regulatory role for glutamine in nephrogenesis. First, an early and dramatic metabolic switch occurs during directed differentiation of both types of hPSCs to NPCs. Second, pathway analysis in both groups revealed major changes in amino acid metabolism, especially in glutamine-related pathways, that was further confirmed in the joint transcriptome-metabolome analysis. Third, deprivation conditions for glutamine enhanced organoid differentiation with changes specifically in podocyte maturation. While broad pharmacologic inhibition of glutamine metabolism interrupted the development of NPCs, specific inhibition of glutamine entry into the TCA cycle with BPTES accelerated the acquisition of advanced NPC morphology, similar to glutamine deprivation. Overall, these results suggest that glutamine catabolism itself plays a critical signaling role in NPC differentiation from pluripotent stem cells.

In recent years, reports on the role of metabolic pathway regulation of renal organoid differentiation have come to light ^24^, and it is clear that metabolic drivers are critically important for differentiation ^25^. Wang, et al, identified the glycine-serine-threonine pathway as providing regulation of proximal tubule development in an embryoid body method of NPC derivation from iPSCs ^26^. Additionally, the idea of supplementation with selected critical metabolites to drive development in a specific direction was demonstrated by the addition of butyrate to enhance proximal tubule development in iPSC-derived renal organoids ^27^. Our study is unique in that it compares and finds commonalities between renal organoid differentiation from hiPSCs and from hESCs and is the first to identify a metabolic driver of podocyte differentiation.

While glutamine is a non-essential amino acid, it plays several key roles in cellular metabolism which make it a key element in metabolic homeostasis, as evidenced by glutamine addiction found in certain cancer cell lines ^23,28,29^. This phenomenon of extracellular nutrients regulating differentiation has been similarly described in other cell types. Glutamine withdrawal was reported to enhance conversion of FOXP3– T cells into FOXP3+ T cells ^30,31^. Additionally, Marsboom et al., suggested that glutamine starvation enhances the angiogenic capability in hESC-derived endothelial cells ^16^. It is clear from the global analysis performed in this study that glutamine metabolism is a key node altered during differentiation of hPSCs to renal organoids.

The mechanisms by which glutamine exerts these effects remain to be discovered. Glutamine metabolism has been associated with OCT4 expression in hESCs ^16^. It is possible that enhanced differentiation under glutamine-deprived conditions may be driven by downregulation of OCT4, expediting the differentiation process ^16^. Glutamine has also been reported to play a pivotal role in promoting dysregulated cell growth in multiple genetic variants of polycystic kidney disease by serving as a precursor to glutathione ^32^. The role of glutamine and other metabolites in promoting dysregulated growth warrants further investigation.

The importance of metabolic regulation in kidney development is highlighted by our findings in the joint analysis of RNA sequencing and metabolomics, which, combined with our metabolomics data on lactate, support a shift from cytosolic glycolysis at the pluripotent stage to mitochondrial oxidative phosphorylation in NPCs. Similarly, Liu et al. reported that self-renewal in murine NPCs requires a high glycolytic flux and glycolysis inhibition promotes differentiation ^33^. Glutamine metabolism has also been connected to podocyte injury, with altered glutamine metabolism in injured podocytes ^34^.

Limitations include that epigenetic factors may still produce disparate results for iPSCs differentiated in the absence of glutamine. Validation with *in vivo* models will shed more light on the extent of glutamine regulation in kidney development. Identifying the specific mechanisms by which glutamine enables this metabolic reprogramming will require further study. The process of directed differentiation from hPSCs to NPCs and renal organoids demonstrates metabolic reprogramming, and our findings further highlight the regulatory role of glutamine in this process. These findings illustrate fundamental features of the developmental biology of the kidney and represent a step towards more efficient renal organoid and tissue engineering.

## Acknowledgements

This research was supported in part by The Hartwell Foundation Biomedical Research Fellowship (I.S.), the Northwestern Medicine Transplant Endowment Grant (I.S. and K.E.H.), the Ruth L. Kirschstein National Research Service Award from the National Heart, Lung, and Blood Institute of the National Institutes of Health under award number F32HL137292 (K.E.H), a grant from the Robert R. McCormick Foundation (J.A.W.), a grant from the Zell Family Foundation (J.A.W.), R01DK113168 (J.A.W.), and Merit Review Award I01BX002660 (J.A.W.) from the United States (U.S.) Department of Veterans Affairs Biomedical Laboratory Research and Development Service. This work has been supported in part by grant UL1 TR001866 from the National Center for Advancing Translational Sciences (NCATS, National Institutes of Health (NIH) Clinical and Translational Science Award (CTSA) program. This research has been supported by the University of Michigan’s George M. O’Brien Kidney Translational Research Core Center, P30DK081943, funded by NIH/NIDDK. This work made use of the (Re)Building a Kidney Coordinating Center funded by the National Institute of Diabetes and Digestive and Kidney Diseases of the National Institutes of Health under award number U01DK107350; the BioCryo facility of Northwestern University’s NUANCE Center, which has received support from the Soft and Hybrid Nanotechnology Experimental (SHyNE) Resource (NSF ECCS-1542205); the MRSEC program (NSF DMR-1720139) at the Materials Research Center; the International Institute for Nanotechnology (IIN); and the State of Illinois, through the IIN. It also made use of the CryoCluster equipment, which received support from the MRI program (NSF DMR-1229693). This work was supported by the Northwestern University RHLCCC Flow Cytometry Facility and a Cancer Center Support Grant (NCI CA060553) and by the Northwestern University Pathology Core Facility and a Cancer Center Support Grant (NCI CA060553). Imaging work was performed at the Northwestern University Center for Advanced Microscopy generously supported by NCI CCSG P30CA060553 awarded to the Robert H Lurie Comprehensive Cancer Center. The iPSC line BJFF.6 was generated by the Genome Engineering and iPSC Center (GEiC) of the Washington University, St. Louis, with the support of the Kidney Translational Research Center (KTRC). The content is solely the responsibility of the authors and do not represent the views of the U.S. Department of Veterans Affairs, the National Institutes of Health or the United States Government.

## Author contributions

IS contributed conceptualization, methodology, validation, formal analysis, investigation, data curation, writing the original draft and reviewing and editing, visualization and funding acquisition. KEH contributed methodology, validation, formal analysis, investigation, data curation, writing the original draft and reviewing and editing the manuscript, visualization and funding acquisition. DI contributed investigation, visualization and reviewing and editing the manuscript. MK contributed formal analysis, data curation, visualization, writing the original draft and reviewing and editing the manuscript. IBS, JMS, MZ, AKG and FHC contributed conceptualization, formal analysis, and reviewing and editing the manuscript. RM and EAO contributed methodology, validation, formal analysis, data curation, visualization and reviewing and editing the manuscript. JAW contributed conceptualization, supervision, project administration, funding acquisition, and reviewing and editing the manuscript.

## Methods

### Contact for Reagent and Resource Sharing

Further information and requests for resources and reagents should be directed to and will be fulfilled by the Lead Contact, Jason Wertheim, MD PhD.

#### Maintenance of H9 cells

The H9 hESC cell line (female, WA09, NIH Approval Number NIHhESC-10-0062) was obtained from WiCell. H9 cells (passage 35–55) and were maintained in mTeSR1 (Stem Cell Technologies, #85850) according to the manufacturer’s protocol. This cell line was authenticated by IDEXX.

#### Maintenance of induced pluripotent stem cells

The BJFF.6 iPSC cell line (male) was obtained from Dr. Sanjay Jian’s lab at Washington University at Saint Louis and were maintained in E8 Flex Medium (Life Technologies #A2858501) according to manufacturer’s recommendations. This cell line was authenticated by IDEXX.

#### Stem cell culture

All hPSCs were passaged using Accutase (Stem Cell Technologies, #07920) or ReLeSR™ (Stem Cell Technologies, # 05872) every 5-7 days. All stem cells were incubated in 10 µM ROCK Inhibitor (Y-27632 dihydrochloride, TOCRIS, #1254) for 1 day after passaging. Routine cytogenetics and mycoplasma testing were performed. All experiments were performed according to applicable guidelines and regulations.

#### Differentiation of hPSCs toward nephron progenitor cells and organoids

Differentiation toward nephron progenitor cells was performed following a modified version of a published protocol by Morizane, *et al*, that was entirely feeder-free ^10^. After a brief wash with PBS, pluripotent stem cells were dissociated with Accutase (Stem Cell Technologies, #07920) or ReLeSR™ (Stem Cell Technologies, #05872). Cells were plated at a density of 40-50 percent coated with 1% Geltrex (Life Technologies, #A1413202) in their respective stem cell media containing ROCK inhibitor Y27632 at 10 µM (TOCRIS, #1254). Approximately 24 hours after seeding, stem cell media was removed, and cells were treated overnight with ReproFF2 supplemented with FGF2 (10 ng/mL). After removing ReproFF2 and washing with PBS, cells were cultured for 4 days in basic differentiation medium composed of 1X L-GlutaMAX (Life Technologies, #35050-061), Advanced RPMI1640 (Life Technologies, #12633-020), and 0.5X pen-strep (Thermo-Fisher 15140122) supplemented with 8 µM CHIR99021 (TOCRIS, #4423). After 4 days, cells were subsequently grown for 3 days in basic differentiation medium supplemented with 10 ng/mL activin A (R&D, #338-AC-050). Media was switched to basic differentiation medium supplemented with 10 ng/mL FGF9 (R&D, #273-F9–025/CF), and cells were incubated for 3 additional days to induce production of NPCs. NPCs were collected at the end of day 9. For continued differentiation to renal organoids, the media was changed to basic differentiation medium supplemented with 10 ng/mL of FGF9 and 3 µM CHIR99021 between day 9 of differentiation to day 11. Media was changed on day 14 to basic differentiation medium (1X L-GlutaMAX in Advanced RPMI1640 with 0.5X pen-strep) and the culture continued for an additional 7 to 14 days (total of 21-23 days). This medium was consistently changed every 2 to 3 days.

#### Immunocytochemistry

Staining was performed as described previously ^10^. Cells were washed in PBS and fixed with 4% paraformaldehyde (Electron Microscopy Sciences, #15710-S) for 15 minutes at room temperature (RT). After three washes with PBS, fixed cells were permeabilized and blocked with 0.3% Triton X-100 and 5% normal donkey serum or 5% goat serum for 1 hour and 30 minutes at RT or with 0.1% Triton X-100 for 15 minutes at RT and Sea block (Thermo Fisher, #37527) for 1 hour at RT.

Cells were subsequently incubated overnight at 4°C with the primary antibody in 1% BSA in PBS with 0.3% Triton X-100. Cells were washed three times in PBS alone and then incubated for 1 hour at RT with secondary antibodies (1:300) (Alexa, Life Technologies). Nuclei were stained with DAPI. Immunofluorescence images were captured on a Zeiss inverted fluorescence microscope or Nikon confocal microscope. Primary antibodies include Oct4 (Life Technologies, #A13998, 1:400), Six2 (Proteintech, #11562-1-AP, 1:100 or 1:300), WT1 (Abcam, #ab89901, 1:200), and Pax8 (Proteintech, #10336-1-AP, 1:1000).

#### Immunohistochemistry on organoids

Organoids were fixed in 4% paraformaldehyde for 1 hour at RT. They were subsequently embedded in HistoGel (Life Technologies, #HG-4000-012) using a Pyrex Cloning Cylinder (Fisher, #09-552-22). Later organoids in HistoGel were paraffin embedded and cut into 4 µm sections. Sections were either stained with hematoxylin & eosin (H&E), or immunochemistry was performed after antigen retrieval. E-cadherin (Life Technologies, #13-1700, 1/100), Podocalyxin (R&D Systems, #AF1658 1/100), Biotinylated Lotus Tetragonolobus Lectin (LTL) (Vector lab, #B-1325, 200 µg/mL).

#### Metabolomics studies

Metabolite extractions were performed as previously described ^35,36^. Briefly, after aspirating medium from the adherent cells in 10 cm dishes, 80% methanol was immediately added to the cells and they were transferred to dry ice and incubated at –80°C for 15 minutes. Then, cells were scraped off the plate on dry ice and the lysate was transferred to 15 mL conical tubes on dry ice and centrifuged at 11,400 g for 5 min at 4°C to pellet cell debris and protein. After washing the pellet with 80% methanol and re-pelleting, the samples were completely dried using a SpeedVac concentrator. Collected samples were transferred on dried ice to the Mass Spectrometry Core at the Beth Israel Deaconess Medical Center for processing following the previously published protocol ^37^. For extracellular metabolomics 1 mL of the collected medium was centrifuged at 14,000 g for 10 min at 4°C. The supernatant was mixed with 4 mL –80°C methanol to make a final 80% (vol/vol) methanol solution. After incubation at 4°C and centrifugation, the samples were dried using SpeedVac and transferred on dry ice for analysis. Metabolites were normalized to the total protein content of the sample.

#### RNA sequencing

Total RNA was collected using the RNAeasy MiniKit (Qiagen) and later was treated with DNA-free™ DNA Removal Kit (Life Technologies, AM1906). RNA QC (quality and quantity) was performed using an Agilent 2100 Bioanalyzer (Agilent). RNA-Seq libraries were generated using Illumina stranded mRNA kits (Illumina). Library QC (quality and quantity) was performed using an Agilent 2100 Bioanalyzer (Agilent). Sequencing was performed on an Illumina HiSeq4000 (single end 50bp) (Illumina).

#### Single cell dissociation of organoid cultures and droplet based single cell RNA-Seq

Organoids were collected by scraping cells from a petri dish (10 cm diameter) and dissociated using cold active protease (*Bacillus licheniformis*) as described by Adam et al., 2017 ^38^. For the 3D organoids, about 20 organoids generated in 3D culture under each condition were dissociated with 5 mg of cold active protease (Sigma, #P5380) in 1 mL DPBS for 15 min on a slow-moving shaker with repeated trituration at 4°C to avoid high temperature artifacts. Cells were passed through a 30 µm cell strainer, washed once in DPBS supplemented with 0.04% BSA, centrifuged at 300 x g for 10 min and adjusted to a concentration of 200,000 cells/mL in 0.04% BSA in DPBS. For the 2D organoid culture, a 10 cm dish containing more than 20 organoids for each condition were utilized. We followed the DropSeq protocol described by Macosko et al. with modifications described in detail by Menon et al ^39,40^. Consistent with our previous findings, in the NPC stage (Figure 5B) ∼500,000 cells were isolated from the standard condition while only ∼200,000 cells were detected under the glutamine-deprived condition. Approximately 100,000 cells of each group were loaded onto a DropSeq device for single cell transcriptomic analysis. Next generation sequencing of generated scRNA-Seq libraries was performed on an Illumina HiSeq2500 platform (asymmetric paired end of 26×115 bp).

#### Flow cytometry

Flow cytometry for PAX8 was performed as previously described, with the following minor modifications ^10^. One million cells were stained with a fixable live-dead stain (Invitrogen, #L34962) and fixed with 4% paraformaldehyde. After permeabilization with 0.1% Triton X-100 for 15 minutes on ice, cells were blocked with PBS containing 1% BSA for 30 minutes on ice and incubated with Pax8 antibody 1:2,500 on ice. The cells were incubated with Alexa Fluor 647-conjugated goat anti-rabbit (Life Technologies, #21244) 1:1,000 on ice. Optimal antibody concentrations were determined using D0 undifferentiated H9 cells and secondary only controls. Flow cytometry was performed on the BD LSR Fortessa cell analyzer. Cells were identified by FSC-A by SSC-A; singlets were identified by FSC-A by FSC-H; live cells were identified by FSC-A by DAPI; and PAX8 positive cells were identified by histogram gated against day 0 H9 cells as well as unstained controls (Figure S5).

#### Cryo-TEM

The High Pressure Frozen (HPF) technique was performed on organoids using a Leica High Pressure Freezer (HPM100). Samples were Freeze Substituted (FS) and processed for resin embedding in the Leica EM AFS2 Freeze Substitution unit. The FS cocktail was 0.5% OSO4, 0.2% UA, in 96% acetone. Samples were transferred under liquid nitrogen into the FS unit, which was set at –90°C. The unit warmed to –80°C over 8 hours, then to 0°C over 16 hours. Three 20-minute acetone wash washes were carried out at 0°C. After the acetone washes, the samples were removed from the FS unit and infiltrated with EMBed 812 resin. Blocks were polymerized in a 60°C oven for 24 hours.

Ultrathin sections of the organoid were produced using a Leica UltraCut S Ultramicrotome, collected on copper mesh grids and post stained with UA and lead citrate. Micrographs were collected using a Gatan Orius camera on a JEOL 1230 TEM and a Gatan Orius camera on a Hitachi HT7700 TEM.

#### Amino-acid deprived ARPMI media

Modified amino-acid-depleted media was generated using RPMI-1640 Medium modified without L-Glutamine, without amino acids, Glucose Culture Media (MyBioSource, #MBS652918). This RPMI1640 was supplemented with amino acids, ascorbic acid (Sigma A4403), sodium bicarbonate (Sigma S5761), zinc sulfate (Sigma Z0251), AlbuMAX II (Thermo Fisher 11021029), insulin-selenium-transferrin (Thermo Fisher 41400045), ammonium metavanadate (Sigma 205559), cupric sulfate (Sigma C8027), manganous chloride (M5005), D-glucose (Sigma G7021), ethanolamine (Sigma E0135) and sodium pyruvate (Sigma S8636) to identically match the published Advanced RPMI1640 formula. Cell survival conditions were verified with custom generated Advanced RPMI1640 purchased from Life Technologies. The custom Advanced RPMI1640 media were used as the base for differentiation media as described above.

### Quantification and Statistical Analysis

All metabolomics samples were collected in triplicate, except D7 for the intracellular metabolomics and D2 for the extracellular metabolomics, for each of which one replicate was lost. The remaining replicates for each day were averaged to generate a third point for analysis. Metabolomics data were analyzed using MetaboAnalyst 4.0 software ^18^.

Flow cytometry data were analyzed with a 2-sided type 2 student’s t-test. Nine replicates, representing separate wells in 6-well plates, were generated for each condition.

For RNA-Seq, raw data was processed into fastq files using Illumina demultiplexing software (Illumina). Raw fastq files were trimmed using TrimGalore (0.4.4) and cutadapt (1.14), with Phred score cutoff of 20 and minimum sequence length cutoff of 20 bp, for removing low quality read adapter sequences ^41^. We then aligned the trimmed fastq files to the human genome sequence (GRCh38.77) using STAR (version 2.5.2b) in quant mode ^42^. The raw counts were then imported into DESEQ2 for differential gene analysis using default parameters ^43^.

For single cell RNA-Seq, FastQC (v0.11.4) was utilized to check the quality of the fastq files from the sequencer, as previously described ^40^. Next, the McCarroll lab (http://mccarrolllab.com/dropseq/) technique was employed, utilizing the embedded Picard tools (picard-tools-1.115) and the DropSeq analysis pipeline to process the fastq files and generate the data matrix table containing the gene expression of the barcoded cells.

Data analysis, correlation analysis, and unsupervised single cell transcriptome clustering was performed as described recently ^40^. Briefly, the downstream analysis consisted of: Normalization, identification of highly variable genes across the single cells, dimensionality reduction (PCA, and t-SNE), standard unsupervised clustering, and the discovery of differentially expressed cell-type specific markers. Combined analysis of cells from organoids derived from glutamine-deprived and standard condition was performed using canonical co-relation analysis function in Seurat R package.

### Data Availability

RNA-Seq datasets generated in this study are available from the NCBI GEO repository under accession number GSE128939. Single cell RNA-Seq datasets generated in this study are available from the NCBI GEO repository under accession number GSE126719. Metabolomics data are available in the (Re)Building a Kidney Consortium collection, https://doi.org/10.25548/16-DYNJ^44^.

## Supplemental Figures

**Figure S1:**
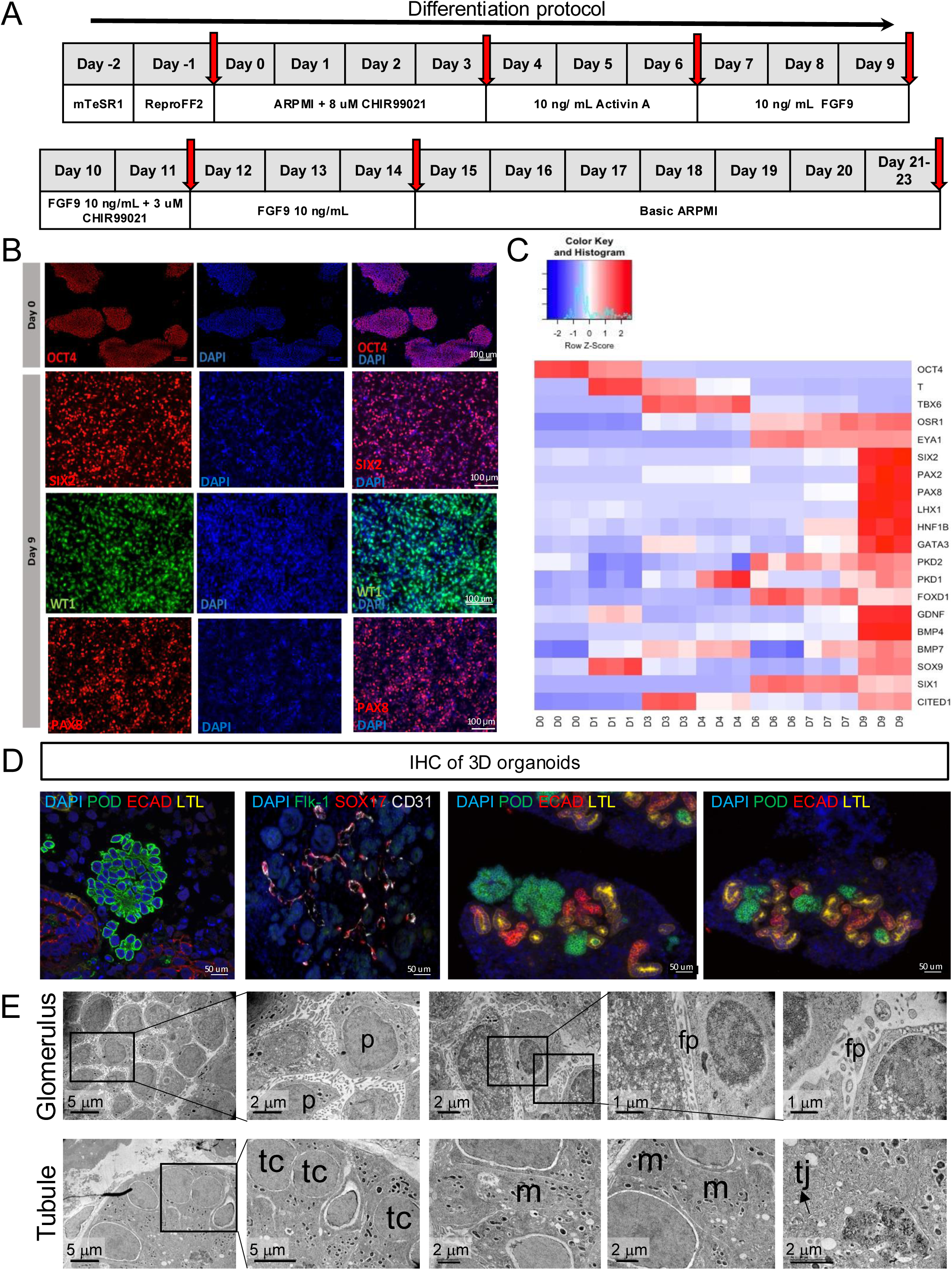
*Nephron progenitor cell (NPC) and renal organoid differentiation.* A) Differentiation protocol to the NPC stage (D9) and organoid stage (D23). B) Immunocytochemistry staining shows differentiation of OCT4 positive H9 cells into SIX2, PAX8 and WT1 positive NPCs at the end of D9. Scale bar 100 µm. C) Heatmap showing dynamic gene expression during differentiation at D0 (day 0), D1, D3, D4, D6, D7, and D9: OCT4 expression decreases in 24 hours after induction, mesoderm markers T and TBX6 expression is detected at D3 and D4, Intermediate mesoderm markers OSR1 and EYA1 is detected at D6 and D7, Nephron progenitor cells and metanephric mesenchyme related genes are detected at D9. Each of three replicates per day is depicted independently. D) Immunostaining depicts generation of renal organoids in 3D culture condition that self-organize into Podocalyxin (POD) expressing early podocytes, E-Cadherin (ECAD) expressing tubular structure and lotus tetragonolobus lectin (LTL) positive renal proximal tubule and SOX17, CD31 positive endothelial cells. Scale bar 50 µm. E) Transmission electron cryo-microscopy (Cryo-TEM) imaging of 3D organoid sections showing podocytes (p), foot process in podocytes (fp), tight-junction (tj), mitochondria (m) and polarized single-layered epithelium of polarized cuboidal tubule cells (tc).

**Figure S2:**
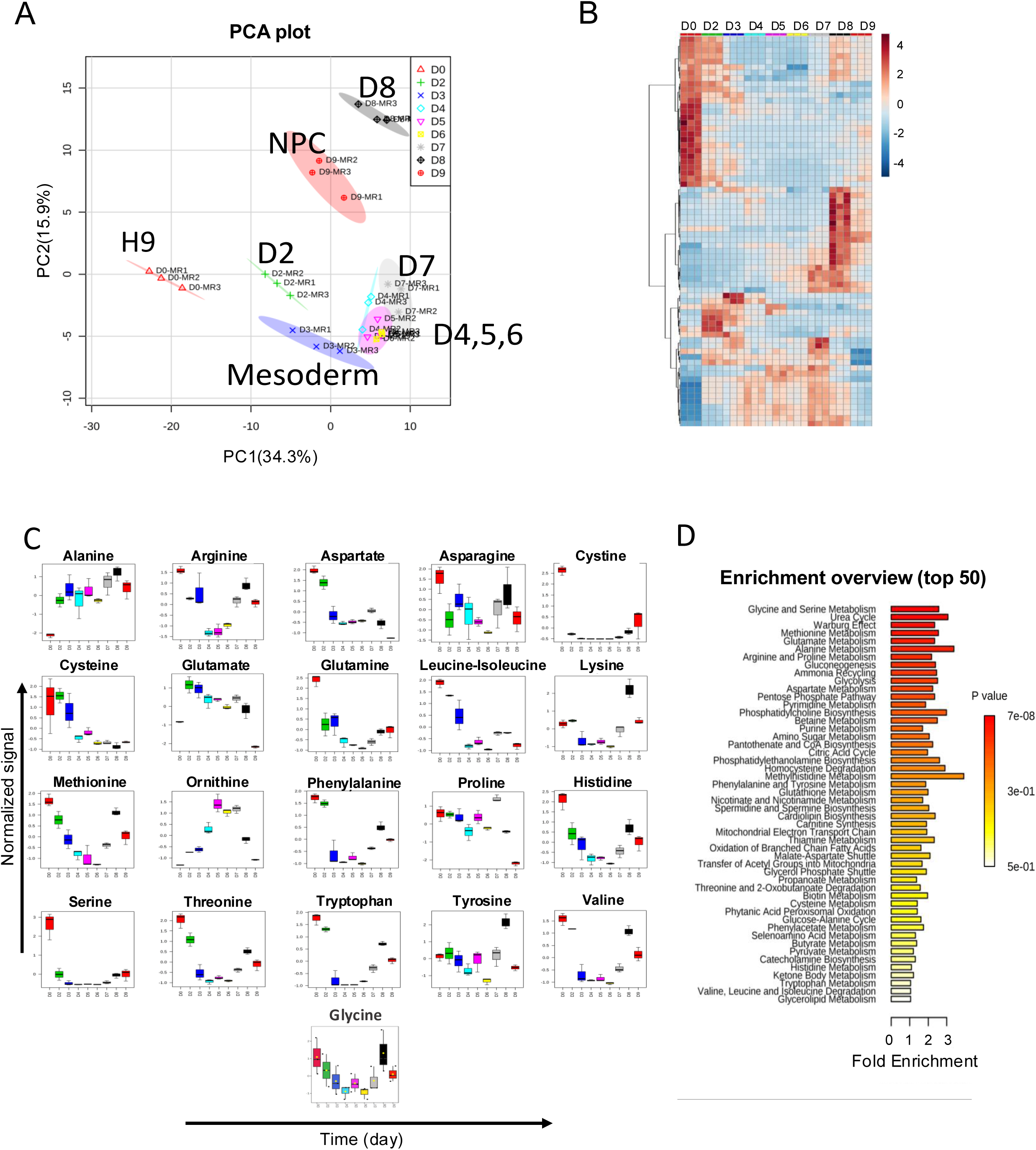
*Extracellular metabolomics of H9 cells to NPCs demonstrate distinct metabolic clusters.* A) Metabolic profiles differ between the mesoderm and NPC stages. Individual dots represent biological replicates. B) Heatmap showing metabolite concentrations (red indicates high, blue indicates low) following three main patterns: Increasing, decreasing and a mid-term maximum at the middle of the differentiation process. C) Changes in the extracellular concentration of amino acids during differentiation. N=3. D) Top pathways in extracellular metabolomics by enrichment analysis.

**Figure S3:**
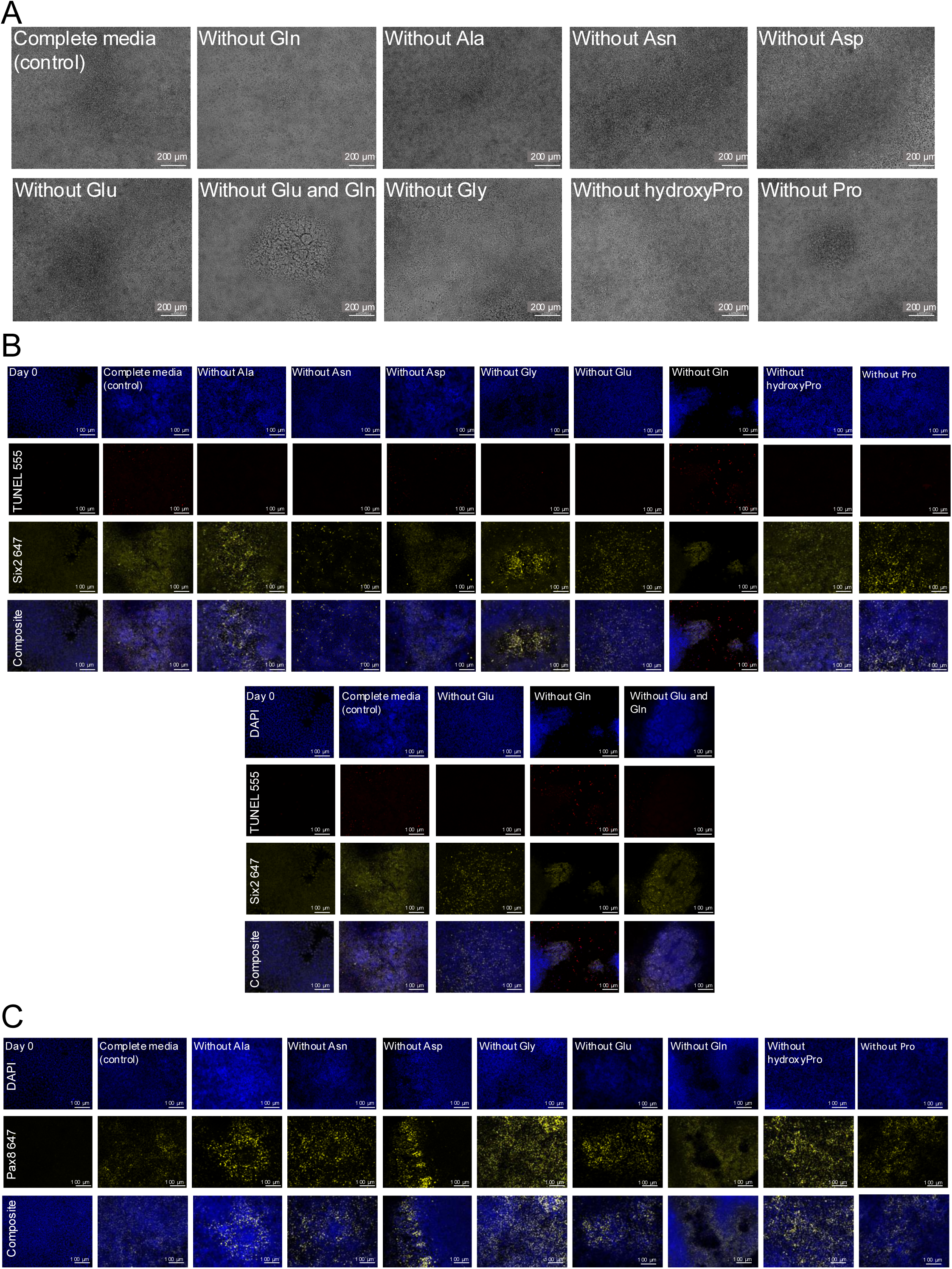
*A subset of amino acids are not required for differentiation to NPCs.* A) All cellular populations that survive single amino acid deprivation demonstrate expected NPC morphology. N=3. B) Cells surviving at the end of D9 in the absence of individual amino acids (Ala, Asn, Asp, Gly, Glu, Gln, Hpro, or Pro) demonstrate SIX2 expression by immunofluorescence. Complete media is used as a control. Day 0, complete media, without Glu or Gln is reproduced, below, to illustrate a comparison to SIX2 expression in the absence of both Glu and Gln. Scale bar 100 µm. C) Surviving conditions demonstrate varying levels of PAX8 expression. Scale bar 100 µm.

**Figure S4:**
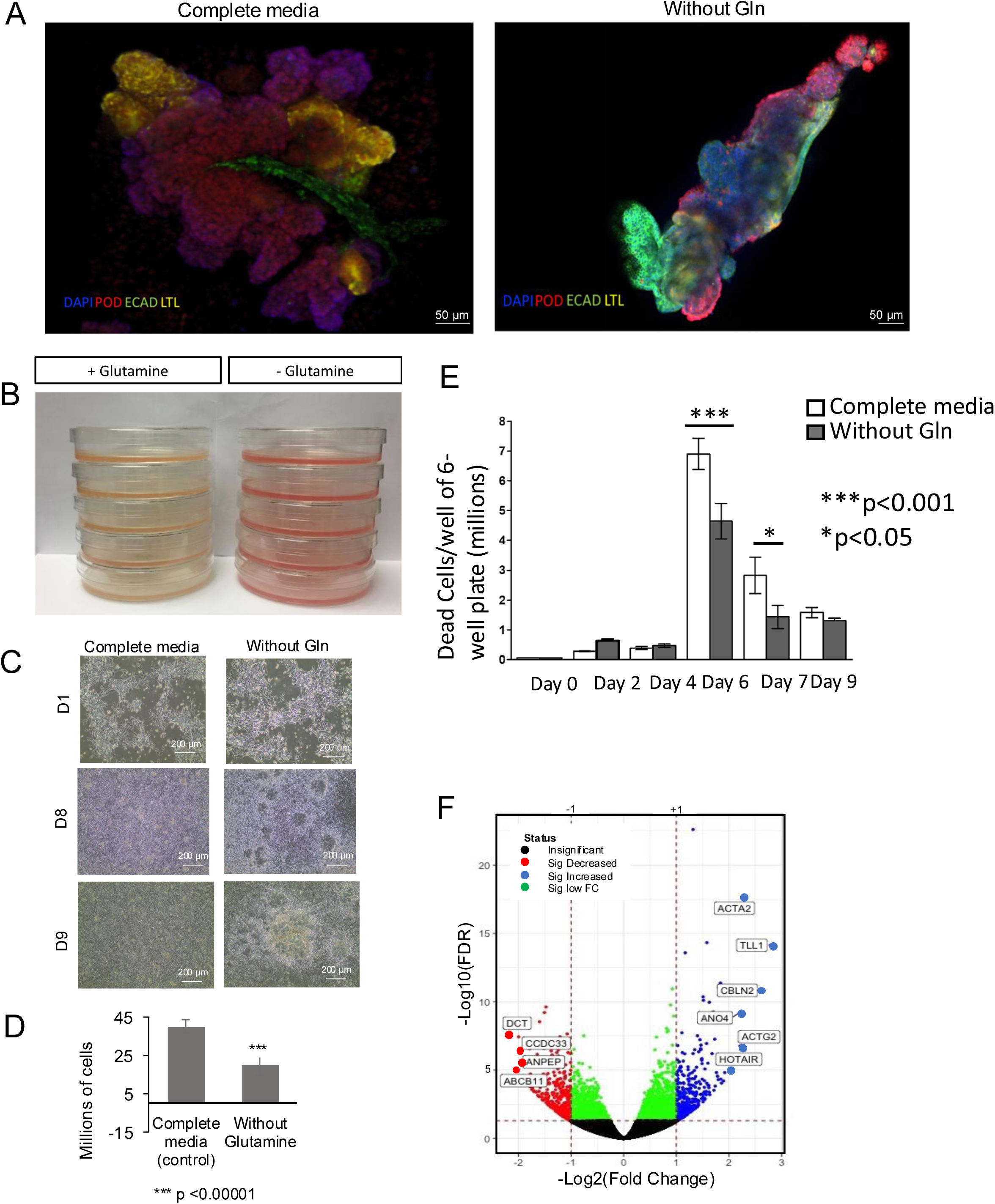
*Total glutamine deprivation yields fewer, but more morphologically advanced, organoids with differential gene expression.* A) NPCs generated in presence (complete media) or absence of glutamine self-organize into renal organoids in 2D culture. Scale bar 50 µm. B) Media color differs in the presence and absence of glutamine, reflecting the lower number of total surviving cells in the absence of glutamine. N=5. C) NPC morphology appears earlier in the absence of glutamine. N=3. D) Number of NPCs recovered in the presence (complete media) or absence of glutamine, mean ± standard error of the mean (SEM). N=3. E) Total glutamine deprivation did not increase cell death. One-way ANOVA. N=3. F) The 10 most highly upregulated genes that differ between NPCs in derived in complete media versus in glutamine deprivation conditions. N=3.

**Figures S5:**
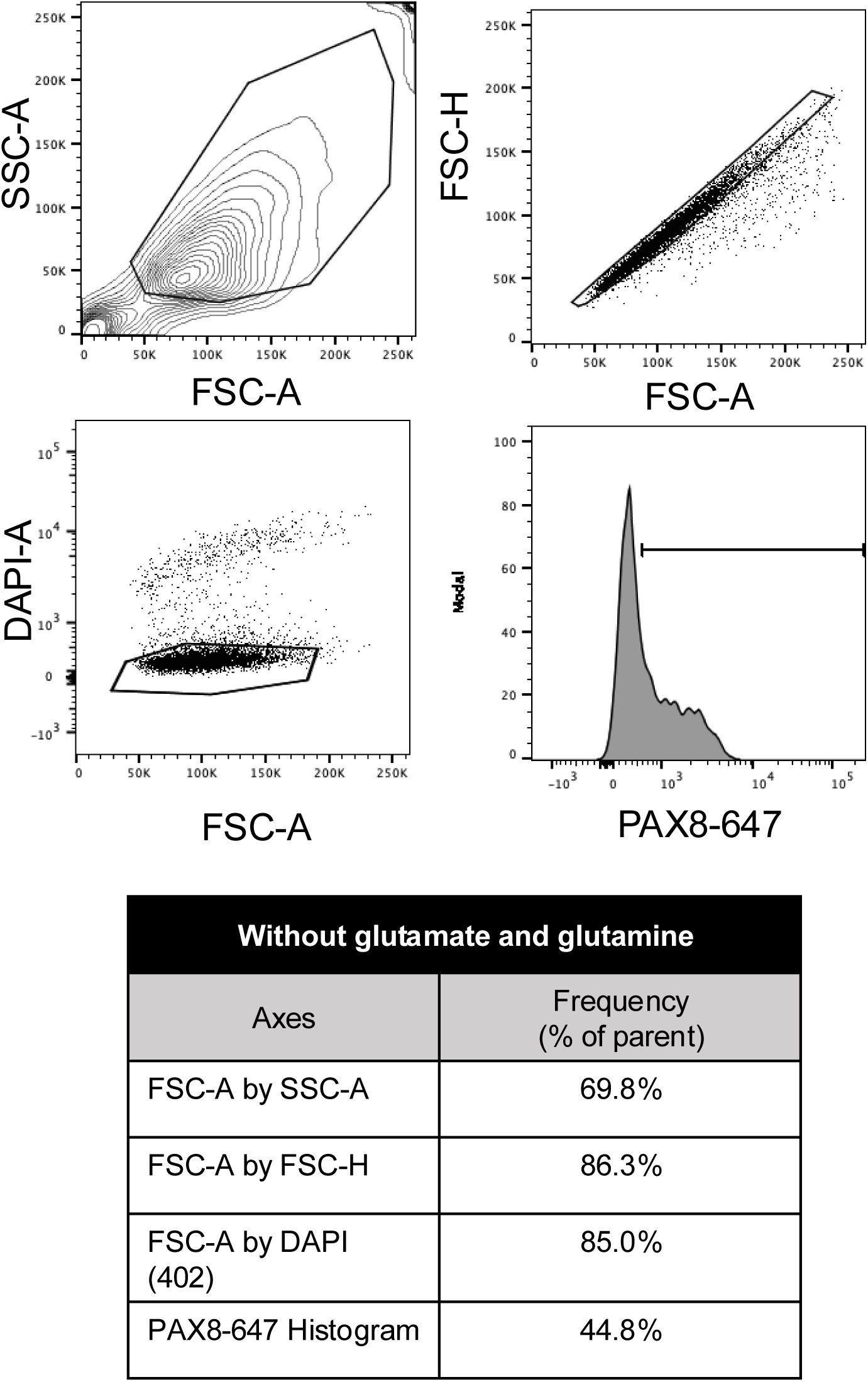
*Sample gating strategy for PAX8 flow cytometry of NPCs grown in the absence of glutamate and glutamine*.

**Figure S6:**
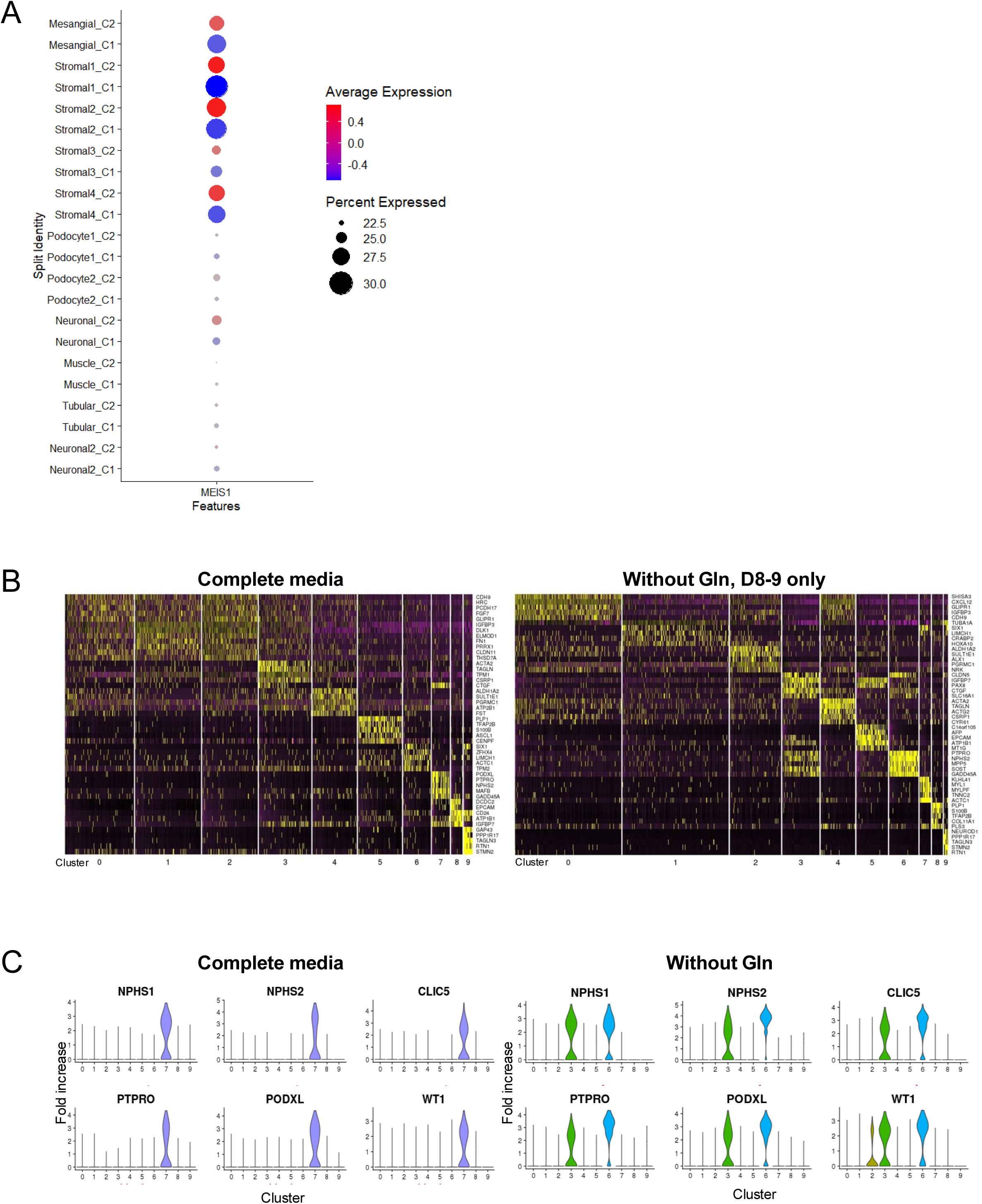
*Glutamine deprivation alters expression of key genes in renal organoids.* A) MEIS2 expression in complete media versus glutamine deprivation. B) Heatmap showing the top differentially expressed genes in each cluster in the presence (complete media) and short-term absence of glutamine. C) Expression of podocyte-related genes in both conditions showing the presence of a second population of podocytes with more robust gene expression.

